# Prior physical synchrony enhances rapport and inter-brain synchronization during subsequent educational communication

**DOI:** 10.1101/601385

**Authors:** Takayuki Nozawa, Kohei Sakaki, Shigeyuki Ikeda, Hyeonjeong Jeong, Shohei Yamazaki, Kelssy Hitomi dos Santos Kawata, Natasha Yuriko dos Santos Kawata, Yukako Sasaki, Kay Kulason, Kanan Hirano, Yoshihiro Miyake, Ryuta Kawashima

## Abstract

Physical synchrony has been suggested to have positive effects on not only concurrent but also subsequent communication, but the underlying neural processes are unclear. Using functional near-infrared spectroscopy (fNIRS) hyperscanning, we tested the effects of preceding physical synchrony on subsequent dyadic teaching-learning communication. Thirty-two pairs of participants performed two experimental sessions. In each session, they underwent a rhythmic arm movement block with synchronous or asynchronous conditions, and then taught/learned unknown words to/from each other according to a given scenario. Neural activities in their medial and left lateral prefrontal cortex (PFC) were measured and inter-brain synchronization (IBS) during the teaching-learning blocks was evaluated. Participants rated their subjective rapport during the teaching-learning blocks, and took a word memory test. The analyses revealed that (1) prior physical synchrony enhanced teacher-learner rapport; (2) prior physical synchrony also enhanced IBS in the lateral PFC; and (3) IBS changes correlated positively with rapport changes. Physical synchrony did however not affect word memory performance. These results suggest that IBS can be useful to measure the effects of social-bonding facilitation activities for educational communication.

## Introduction

Physical synchrony is an important nonverbal factor in forming and modulating communications^1^. Many studies have reported generally positive effects of physical synchrony on affective and cognitive aspects in communications^2–4^. Experimental manipulation of physical synchrony induced higher affiliation^5^, cooperation^6^, compassion and altruistic behaviour^7^, and coordination performance^8^, better mood^9^, and incidental memories^10^. Since successful teacher-student communication is a key element of education, the effect of physical movement synchrony between teacher and student during lecture has been investigated and associated with higher rapport^11, 12^ and better learning operationalized by higher student quiz scores^13^.

Furthermore, physical synchrony can have not only concurrent but also prolonged effects on subsequent social interactions. Physical synchrony prior to a task has been shown to lead to enhanced cooperative behaviours between children^14^. Synchrony of body and head movements between therapists and patients predicts short- and long-term psychotherapy outcomes^15^. These results suggest the effectiveness of various practices involving physical synchrony for the enhancement of communication. Indeed, practitioners and researchers in the field of education have recognized the effect of getting “in synch” with students, and have incorporated synchronous components such as singing and dancing as “warming-up” activities to facilitate better social and cognitive outcomes in the classroom^16–18^. However, psychological and neural processes underlying the sustained effects of physical synchrony need further exploration.

Recent studies measuring multiple brains simultaneously, termed “hyperscanning”, have revealed that various aspects of communication can be reflected in the interpersonal neural synchronization or inter-brain synchronization (IBS) between the individuals engaging in communications^19–24^. IBS is a measure of functional connectivity between the different individuals’ regional neural activity changes, reflecting how much their brains are recruited in a temporally coordinated manner. It has been shown that cooperative communication or interaction^25–29^ and availability of communication-facilitating nonverbal factors in face-to-face situations^30, 31^ generally enhance IBS in the communication task-related brain regions such as the prefrontal cortex (PFC) and the temporo-parietal regions. IBS has also been reported to reflect the efficiency of communications^32–36^ or the strength of the bond with a peer^37–40^. These results suggest that the application of hyperscanning to the evaluation of IBS in the field of educational communication is promising^41–46^, especially to probe into the degree of harmonization in social affective and cognitive processes involved in the communication. However, although several studies reported changes in IBS during coordinated physical activities^47–50^, no study so far has directly addressed the link between the positive effect of prior physical synchrony and IBS during subsequent communication.

The main objective of this study was thus to investigate the effects of prior physical synchrony on subsequent educational communication as well as prefrontal IBS during the communication. We hypothesized that (1) prior physical synchrony has positive psychological carry-over effects on the communication; (2) it also leads to higher prefrontal IBS during the communication; and (3) the effects on the psychological outcomes of communication and IBS are positively associated. To test these hypotheses, we conducted an experiment consisting of four steps per session (Fig. 1a). Pairs of participants, provided with a story claiming that the study purpose was “to investigate the effect of embodied physical tempo on the succeeding learning process”, experienced incidental synchronous or asynchronous physical movement, Then, they engaged in an educational communication where they taught and learned unknown words to/from each other. Their neural activities in the medial and left lateral PFC, which are involved in internally and externally oriented cognitive functions, respectively^51–53^, and whose activities have been shown to synchronize interpersonally during verbal communication^26, 33, 36^, were measured using a two-channel portable wireless functional near-infrared spectroscopy (fNIRS) device (Fig. 1b, c). fNIRS utilizes the link between changes in neuronal activities and cerebral blood flow and chromophore concentration, a mechanism known as neurovascular coupling^54, 55^. fNIRS has the advantages of relatively high cost-effectiveness, tolerance to motion artefacts, and the localized nature of hemodynamic effects^55^, which make it a promising application for education and other real-world communication. After finishing the teaching-learning communication task in each session, participants rated the rapport they felt during the task, and completed a memory test about the taught/learned contents.

**Figure 1.**
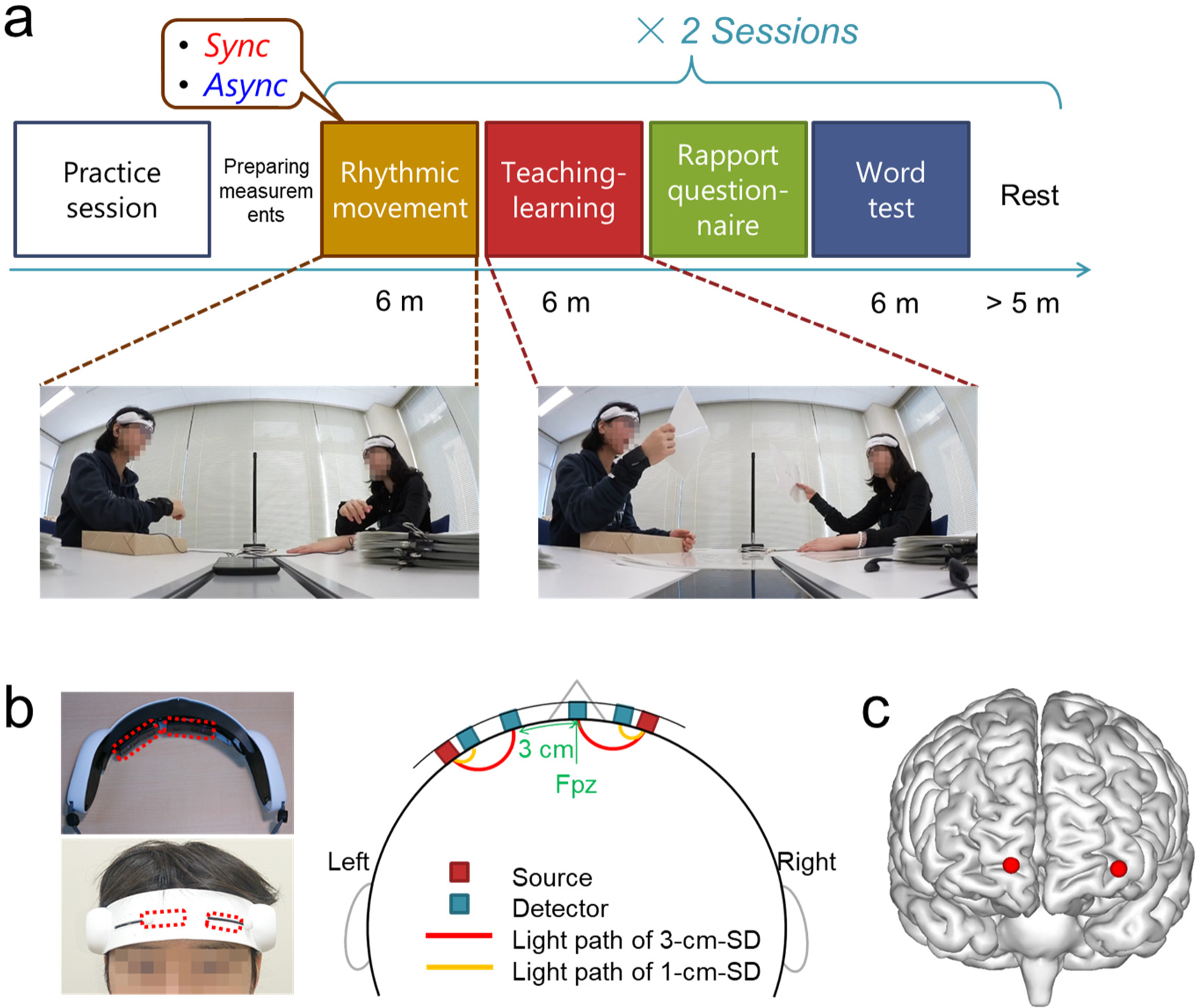
Experimental procedure and fNIRS measurement setup. (a) Flow of the experiment, with snapshots of the rhythmic movement (Sync, synchronous; Asynch, asynchronous arm movements) and the word teaching-learning blocks. (b) Configuration of the fNIRS optodes, channels, and their placement on the prefrontal region. See Methods for details on the deep (3-cm-SD) and shallow (1-cm-SD) channels. (c) fNIRS channel positions projected on the normalized Montreal Neurological Institute (MNI) brain template, estimated using the virtual registration^77^. The red spheres indicate the 3-cm-SD channels, which are located in the middle between the light sources and the 3-cm detectors. The visualization was created using BrainNet Viewer^91^.

## Results

### Effects of prior physical synchrony on rapport and memory outcomes

In order to test our first hypothesis that participants achieve a stronger mutual bond in the teaching-learning task after experiencing synchronous physical movement (post-Sync) than after asynchronous physical movement (post-Async), we conducted a paired t-test on the averaged pair-wise ratings on the rapport they felt during the teaching-learning task. We found a significant effect of prior physical synchrony on rapport (*t*(31) = 3.55, false discovery rate (FDR)-adjusted *p*_FDR_ = 0.001; Fig. 2). As a post-hoc analysis, we also tested the effect separately on the three components of rapport: mutual attentiveness, positivity, and coordination^56^. All components showed significantly higher scores in the post-Sync than the post-Async condition, with the coordination component showing the strongest effect (mutual attentiveness: *t*(31) = 1.91, *p*_FDR_ = 0.036; positivity: *t*(31) = 1.86, *p*_FDR_ = 0.036; coordination: *t*(31) = 3.99, *p*_FDR_ < 0.001; Supplementary Fig. S1a-c).

**Figure 2.**
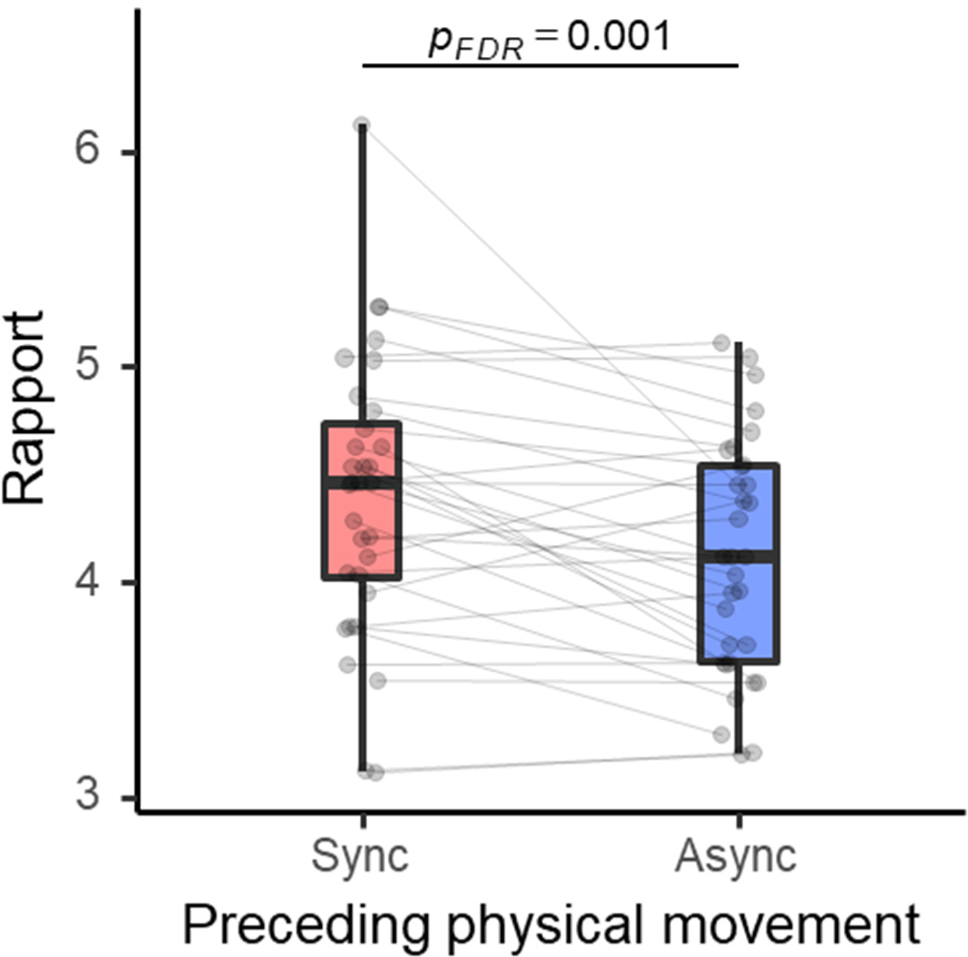
Subjective rapport during the word teaching-learning task after experiencing physical synchrony/asynchrony. Box plots show the median, interquartile range (IQR), and minimum/maximum values of the average rapport ratings during the teaching-learning block of all dyads for the two conditions of the preceding rhythmic movement block (Sync, synchronous; Async, asynchronous arm movements). Grey points with connecting lines represent the two conditions of each dyad.

Furthermore, to test the possibility that the enhanced rapport might also transfer to better focus during the communication and thus result in better memory encoding of the taught and learnt contents, we conducted a paired t-test on the averaged pair-wise memory test scores. We observed no significant effects of post-Sync vs. post-Async conditions (*t*(31) = −0.74, *p*_FDR_ > 0.05; Fig. 3a). Additionally, participants’ subjective ratings of the effectiveness of learning did not show significant effects of prior synchronous/asynchronous physical movement (*t*(31) = 0.22, FDR *p*_FDR_ > 0.05; Supplementary Fig. S2). As a supplementary analysis to investigate the potential influence of the roles of the participants (teacher/student) and a possible interaction effect with the prior physical synchrony/asynchrony on the word memorization process, we calculated the average word test score separately for words that a participant experienced as a teacher and words that he/she experienced as a student (Fig. 3b). We then conducted a two-way repeated measures analysis of variance (ANOVA). A significant main effect of role was observed (*F*(1,31) = 8.72, *p* = 0.006), where word memory as a teacher was poorer than word memory as a student. Neither the interaction (*F*(1,31) = 0.38, *p* = 0.540) nor the main effect of the post-Sync/Async condition (*F*(1,31) = 0.69, *p* = 0.412) were significant.

**Figure 3.**
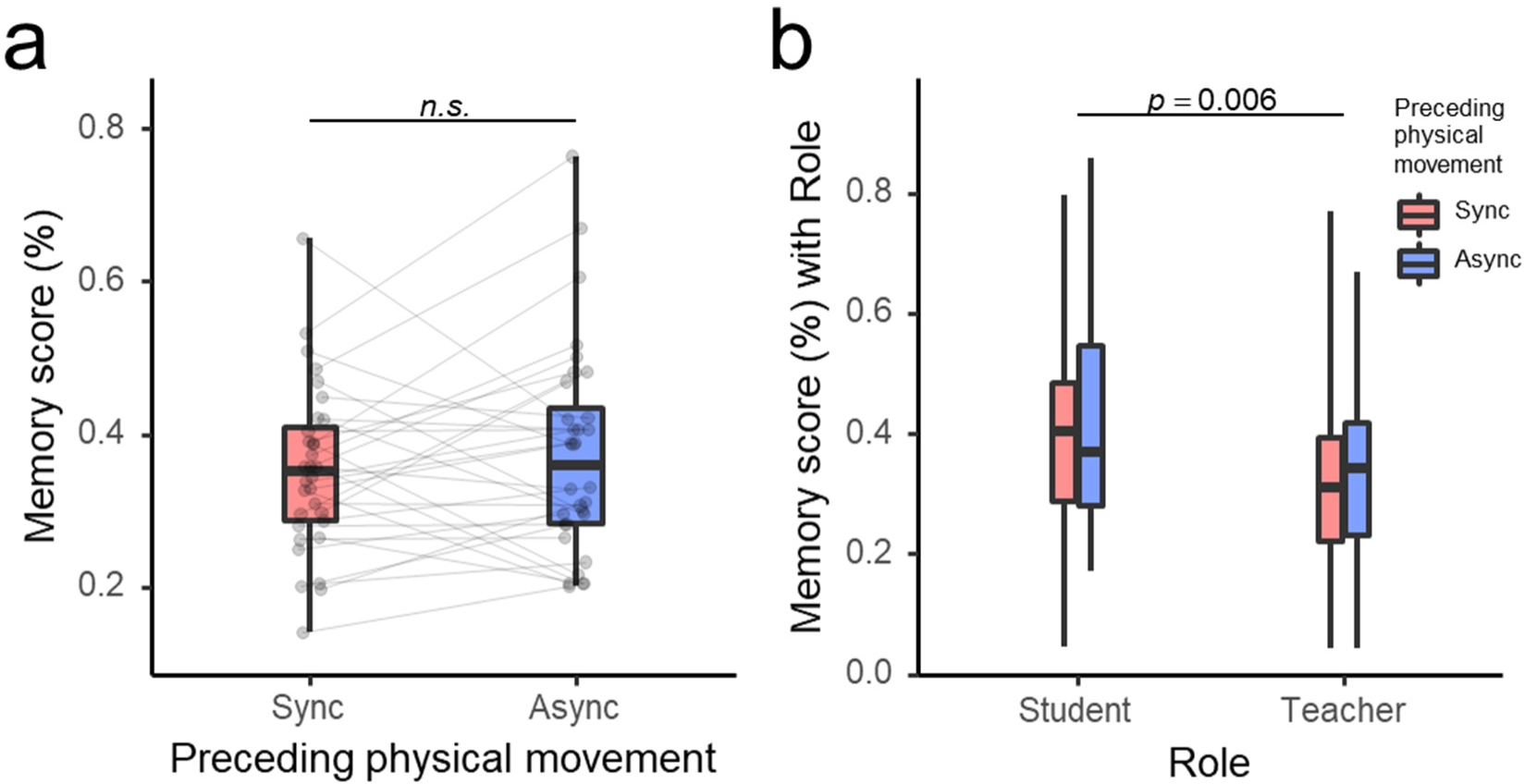
Word memory test scores after experiencing physical synchrony (Sync)/asynchrony(Async). (a) Total scores. (b) Scores separately for words experienced as a teacher and as a student. See the caption of Figure 2 for box plot notations.

### Prior physical synchrony enhanced IBS during teaching-learning task

IBS between the teaching and learning pairs’ homologous activities in the left lateral and medial PFC was evaluated using wavelet transform coherence (WTC)^57^ (Supplementary Fig. S3). The IBS values were calculated by averaging the WTC *R*^2^ at the Fourier period corresponding to the task cycle (45 s) over the whole teaching-learning block. Paired t-tests were conducted to test our second hypothesis that IBS during the teaching-learning task was higher in post-Sync than in post-Async conditions. The results showed significant effects of prior physical synchrony on the IBS in the left lateral PFC (*t*(31) = 3.49, *p*_FDR_ = 0.001; Fig. 4a), but not on the IBS in the medial PFC (*t*(31) = 1.51, *p*_FDR_ > 0.05; Fig. 4b).

**Figure 4.**
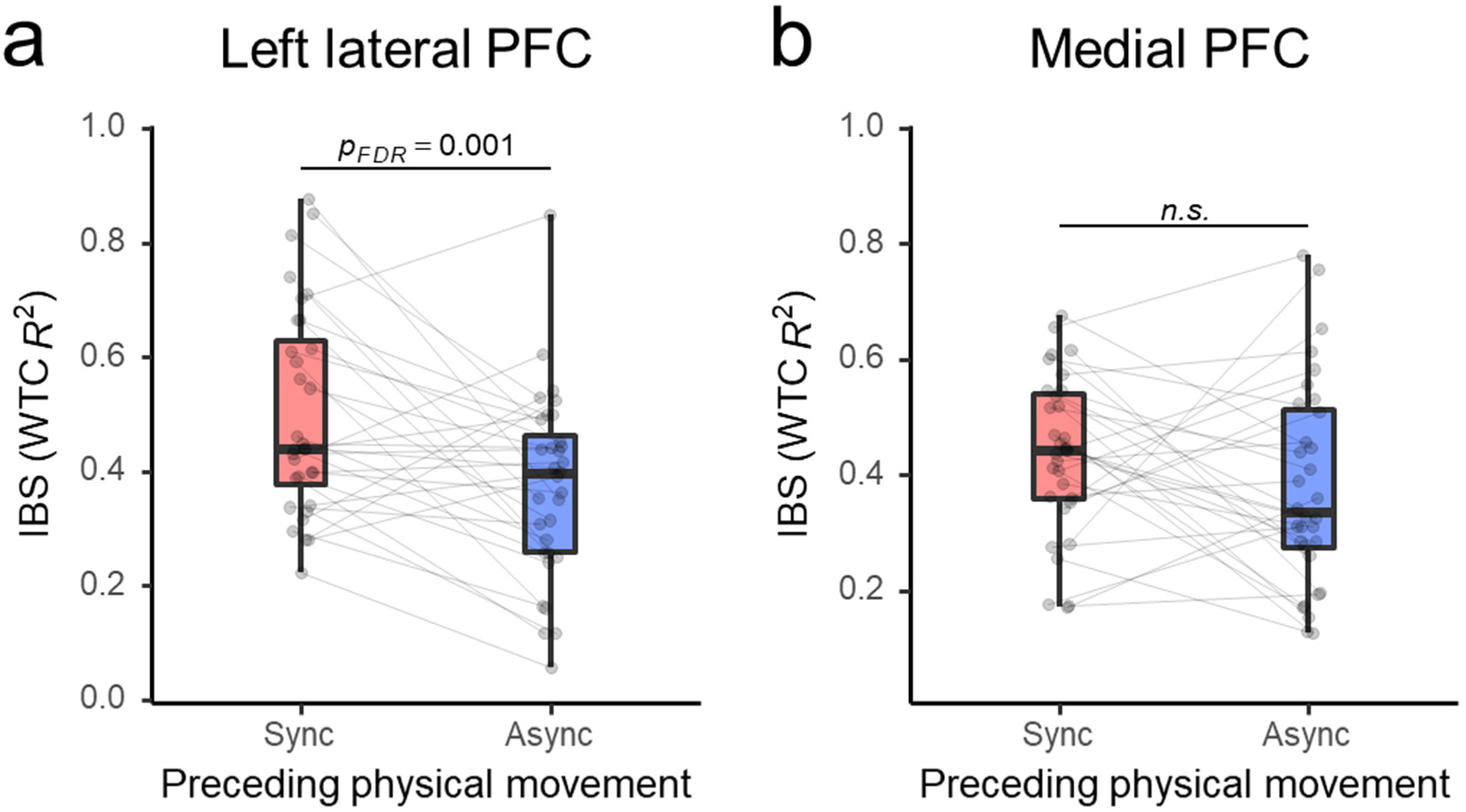
Inter-brain synchronization (IBS) in the left lateral prefrontal cortex (PFC; a) and the medial PFC (b) during the word teaching-learning task after experiencing physical synchrony/asynchrony. See the caption of Figure 2 for box plot notations. WTC, wavelet transform coherence (see main text for details).

In order to confirm the origin of the observed effect on IBS, we also evaluated the interpersonal synchronization of the “shallow signals” from the adjacent short source-detector (1-cm-SD) channels, which mostly reflect blood flow changes in the tissues shallower than the brain^58, 59^. They showed no enhancing effects, neither in the left lateral nor the medial channels (*t*(31) = −0.27 and *t*(31) = −0.28 respectively, both *p*_FDR_ > 0.05; Supplementary Fig. S4a, b). Furthermore, we also compared the interpersonal synchronization of the fNIRS signals that were not corrected for the shallow signal component (see Methods for details). Again, we observed no significant difference between the conditions in either channel (left lateral: *t*(31) = 1.11, medial: *t*(31) = 1.98, both *p*_FDR_ > 0.05; Supplementary Fig. S4c, d).

### IBS and rapport were positively correlated

To test our third hypothesis of a positive relationship between the degrees of rapport and the IBS in the left lateral PFC, which both showed enhancement induced by prior physical synchrony, we evaluated the within-subject correlation between these variables, and found a significant positive correlation (repeated measures *r*_rm_(31) = 0.42, *p*_FDR_ = 0.010; Fig. 5). The results indicate that the effects of prior physical synchrony manifested in these two measures of teaching-learning communication in a positively associated manner. As a post-hoc analysis, we tested the correlation between the left lateral PFC IBS and the three components of rapport. Mutual attentiveness and coordination showed significant positive within-subject correlation with IBS, while positivity did not show a significant correlation (Mutual attentiveness: *r*_rm_(31) = 0.34, *p*_FDR_ = 0.038; positivity *r*_rm_(31) = 0.21, *p*_FDR_ > 0.05; coordination *r*_rm_(31) = 0.43, *p*_FDR_ = 0.020; Fig.S3a-c).

**Figure 5.**
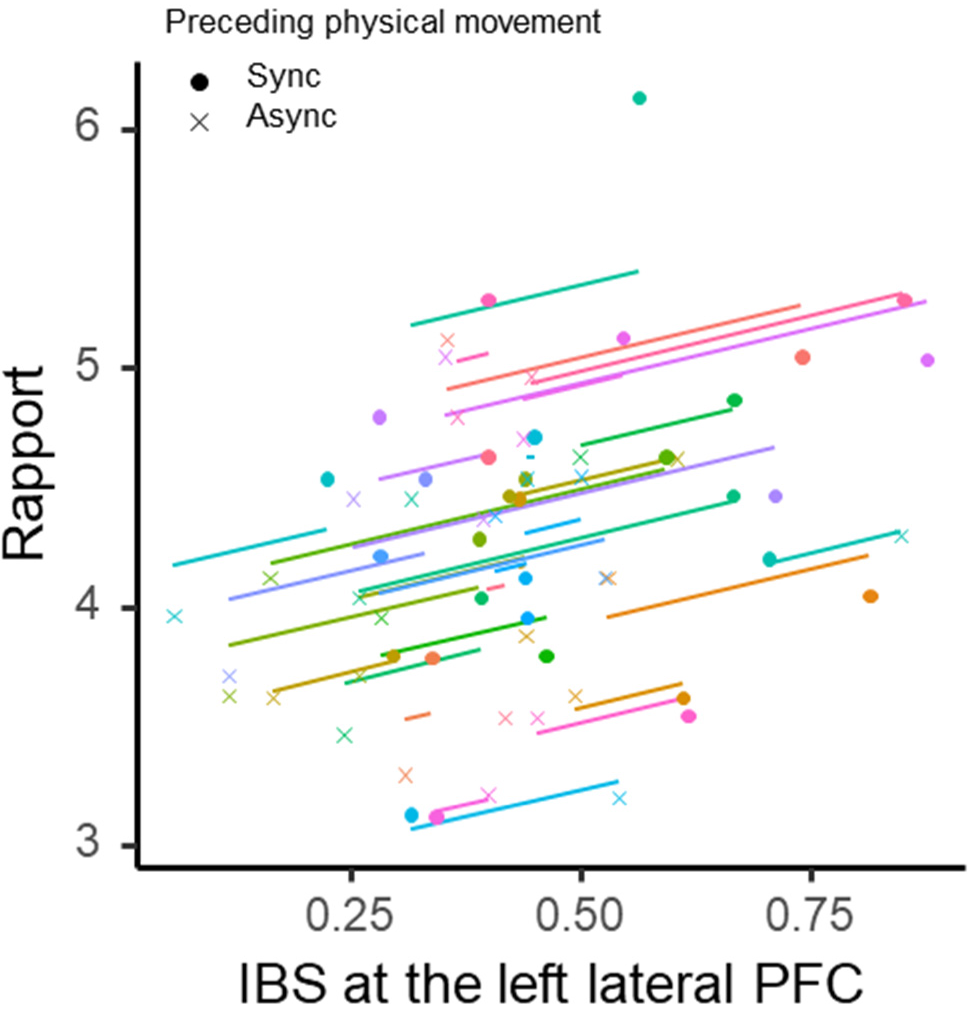
Within-pair correlation between inter-brain synchronization (IBS) at the left lateral prefrontal cortex (PFC) and the rapport ratings during the word teaching-learning task modulated by prior synchronous (Sync) and asynchronous (Async) physical movements. Coloured points (•, ×) represent the two conditions of each dyad. Coloured lines represent the best linear fit for each dyad estimated by the repeated measures correlation, using the same slope and varying intercepts^89^.

## Discussion

The current results generally support our three initial hypotheses: (1) prior physical synchrony enhanced rapport between teacher and learner; (2) prior physical synchrony enhanced IBS in the left lateral PFC during the teaching-learning task; and (3) the effects on the rapport and left lateral PFC IBS were positively correlated. In the following, we discuss possible mechanisms and implications of these results, relationships with previous studies, and limitations of the current study.

Our analysis confirmed positive effects of preceding physical synchrony on the social bonding perceived as rapport during the subsequent communication. This observation extends previous findings that physically synchronous behaviours between teacher and student during lecture is linked to higher rapport^11–13^. The result is also in line with a previous report that passive experience of physical synchrony prior to a task leads to enhanced cooperation between children who had been unfamiliar to each other before the experiment^14^. This indicates that prior physical synchrony is indeed a useful “warming-up” method to achieve higher levels of bonding during communication.

An interesting question here would be to which degree the effect was mediated by conscious/top-down or unconscious/bottom-up processes. Previous psychological studies have provided indecisive evidence for the roles of awareness and intentionality factors^3, 4^. Although this study cannot provide a resolute conclusion on this issue, (a) the fact that participants confirmed to have been unaware of the true hypothesis in the debriefing after the experiment, and (b) the observation that among the three components of rapport, coordination, with cues that are complex and less available to conscious control than positivity and attentiveness cues^56^, showed the greatest enhancing effect, suggest that the prior physical synchrony effect on the subsequent communication was mediated primarily via a bottom-up pathway.

On the other hand, we did not observe an enhancing effect of prior physical synchrony on word memory scores. This is in contrast with a previous study where a teacher’s imitative behaviours of students during lectures led to enhancement in both rapport and memory performance^13^. There are several possible explanations for the seeming discrepancy. A main factor is differences in the timing and types of coordination. In the previous study^13^, an experienced confederate teacher was allowed to imitate the students’ wide range of nonverbal behaviours during the lecture. Since the imitation was directly embedded in the teaching activity, it may have been effectively used to manipulate the students’ attention toward appropriate aspects at the appropriate timing. In this study, on the other hand, physical synchrony in a restricted form was experienced prior to the teaching-learning task, and the effects thus lack any specificity regarding aspects and timing relevant to the cognitive states of teacher and learner or the contents being communicated. Another (somewhat related) reason for the discrepancy might stem from the design and contents of the educational communication task used in this study. Participants who did not have particular experience in teaching took turns playing the role of teacher or student, and adhered to the restricted teaching-learning procedure that excluded any kind of improvisation (see Methods). Therefore, independently of the teacher’s effort, on the student’s side it might have been a more efficient strategy for good memory encoding to grasp the needed information (word meaning and example sentence) at the beginning and recite it internally while ignoring the teacher in the later phase of each trial. Furthermore, in contrast to the previous study^13^, participants were also required to memorize the new information while teaching it. These factors may have led to a trade-off or resource competition between the social cognition directed to the partner, which is linked to rapport, and the information processing linked to memorization. The possibility of such resource competition occurring here is supported by the supplementary observation that memory for words experienced as a teacher was significantly poorer than memory for words experienced as a student. Interestingly, in a previous study that investigated the effect of physical synchrony on incidental memories, subjects picked up distracting word utterances from the synchronizing partner more than from the non-synchronizing partner, even though they were instructed to ignore those word utterances^10^. All these findings lead us to speculate that memory performance might also have been enhanced by prior physical synchrony, if the encoding target was related to or generated by the partners on the spot, where the attention to and bonding with the partner would have been more beneficial, or in a more adaptive setting where the teachers were allowed to devote themselves to the teaching and to adjust the style and contents flexibly.

As a side remark, we would like to note that the observed non-effect of the preceding physical synchrony on memory scores in the subsequent learning task at one shot should not be generalized to its uselessness for cognitive outcomes in real education, even if the teaching is provided in a similar manner as in the current experiment. An important factor to be taken into consideration would be the learner’s motivation. In fact, it is still possible that enhanced rapport induced by physical synchrony might lead to a better attitude toward learning, thereby leading to better cognitive outcomes in the long-term.

Our experiment revealed that neural activities in the left lateral PFC of the pairs engaging in the teaching-learning task were significantly more synchronized after synchronous arm movements than after asynchronous movements. This finding of prior physical synchrony enhancing IBS in the subsequent communication may be regarded as the most significant contribution of the current study. Previous physical coordination-related IBS studies observed interpersonal neural coordination *during* physical interaction^47–50^. Moreover, there is an fMRI study that probed into individual brain activity changes during and immediately after synchronous/asynchronous beat generation with ostensible partners (actually a computer program)^60^. However, to the authors’ knowledge, no study so far has shown effects of preceding physical synchrony on interpersonally coordinated brain recruitment during subsequent communication.

Previous studies have indicated functional and structural differences between the lateral and medial parts of the anterior prefrontal region^53, 61, 62^, with the lateral part responding more to highly demanding and often externally-oriented cognitive domains and being functionally and anatomically connected to the “task-positive networks” such as the executive control and salience networks, whereas the medial part responding more to low-demand and internally-oriented situations and being connected to other components of the default mode network^52, 63, 64^. Thus, our results suggest that prior physical synchrony enhanced the coordinated recruitment and/or regulation of externally oriented cognition. More specifically, the lateral anterior PFC has been implicated in working memory, retention and switching of multiple task sets, and in sustained cognitive control^53, 65, 66^, all of which would have been important for the participants’ engagement in the teaching-learning task.

On the other hand, IBS in the medial PFC, which has been implicated in internally-oriented affective and cognitive processes, especially social mentalizing (i.e. understanding the beliefs, emotions, desires, and dispositions of others)^51, 52, 67^, did not show significant enhancement induced by prior physical synchrony. This may be attributable to the asymmetric nature of the teaching-learning communication. Unlike during daily communication where people take turns speaking in an unconstrained and bidirectional manner^30, 26^, during teaching-learning communication, which aims to transmit knowledge, a teacher needs to infer the student’s knowledge and current state of understanding, while the student can focus on the transmitted information itself and is less required to think about the teacher’s mental states^46^. This might have led to the asymmetrical recruitment of mentalizing functions, and resulted in no effects of prior physical synchrony on IBS in the medial PFC. Based on these speculations, one could expect that more person-oriented (mutually attentive) rather than knowledge-oriented communication, such as collaborative learning aiming at exchanging and cooperatively building ideas, would require more mentalization in both the speaker and listener^68^, and could thus lead to enhanced IBS induced by prior physical synchrony also in the medial PFC. This is an interesting possibility to be explored in future studies.

The current results are unlikely to be attributable to modulated synchronization of artefactual signals originated from autonomic activities or physical head motions during the teaching-learning block, as the supplementary analyses showed no effect of prior physical synchrony on the interpersonal synchronization between the shallow tissue signals, which do not contain neural signals and are dominated by autonomic and physical components (Supplementary Fig. S4a, b). Additionally, a more significant effect on IBS was detected after the removal of shallow signal (Fig. 3) in comparison to the typical fNIRS signal (3-cm-SD) without removal (Supplementary Fig. S4c, d).

These results indicate that the observed effects of the prior synchronous vs. asynchronous physical motion was indeed of neural origin. They are also consistent with the previous studies^26, 50^ that showed the effectiveness of utilizing referential shallow signals for artefact reduction and sensitive IBS detection in natural interaction settings.

Our finding of a positive association between the modulated rapport and IBS in the left lateral PFC suggests that the coordinated recruitment of the lateral prefrontal region, induced by preceding physical synchrony, may be the underpinning of the enhanced rapport. This is also in line with the previous studies that suggested that coordinated activities in the left lateral PFC between teachers and students are associated with their successful interaction^42, 43^. Furthermore, looking into the three components of rapport, IBS in the left lateral PFC was significantly correlated with mutual attentiveness and coordination, but not with positivity. Considering that attentiveness and coordination are perceptually guided (i.e. externally oriented) judgments, while positivity represents an affective inference (i.e. internally oriented) judgment, these results are consistent with the interpretation that enhanced IBS in the lateral PFC reflects harmonized cognitive engagement toward the partner or toward the shared activity at hand.

A major limitation of the current study is the regional coverage of the fNIRS measurement. Unfortunately, the portable fNIRS device used in this study allowed the measurement of only the regions covered by hairless scalp skin. Future studies using a device that provides wider coverage of regions, especially the language-related and additional social-related regions, such as the inferior frontal gyrus and temporo-parietal regions focused on in previous IBS studies^30, 69, 31, 70, 36, 46^, could shed further light on IBS in those regions. Open questions are, for example, how IBS in these regions could reflect verbal or perceptually-guided social processes (e.g. the learner mentally shadowing utterances of the teacher or reading states/intentions from the partner’s nonverbal cues), how such processes could be affected by prior physical synchrony, and how they could lead to changes in learning performance. On the other hand, to those who intend to apply fNIRS brain hyperscanning to the objective evaluation of communication and its enhancement in educational and other real-world contexts, it will be encouraging to see that hyperscanning using such (relatively) low-cost and portable devices can successfully capture such hypothesized effects. Furthermore, the auxiliary measurement of shallow signals and their use in the removal of artefacts is a strength of the current study in confirming the neural origin of the observed IBS effects.

Physical movement synchronization could have been introduced in different manners in regard to various factors, such as phase (in-phase vs. anti-phase), positioning of the partners (face-to-face vs. side-by-side), awareness and intentionality regarding the synchronization, etc. Although differential psychological effects of these factors have been studied to some extent^3^, future study is required to determine whether such differential effects involve IBS changes in different regions (reflecting different mechanisms) or are merely related to different degrees of IBS in the same neural region. A specific avenue that warrants being pursued further would be the role of metacognition of the physical synchrony in the subsequent communication. As discussed above, our experimental design and results suggest a relatively minor impact of metacognition on average. Nevertheless, at the individual level, it is possible that some dyads may have been more self-aware of the physical synchrony than others, and such differences might have led to either increased or decreased effects on the following teaching-learning processes. The issue of whether to encourage or inhibit metacognition of the physical synchrony will also be relevant to the successful design of communication-enhancing activities in educational and other practical fields.

In conclusion, this is the first study that verifies the associations between preceding physical synchrony and psychological beneficial effects in association with IBS during the subsequent communication. The results suggest the usefulness of brain hyperscanning in analysing the effects of various “warming-up” activities that are used as daily practices in education and other communication fields to facilitate social bonding.

## Methods

### Participants

Sixty-four healthy, right-handed Japanese undergraduate and graduate students participated in this study (age range: 20−26, mean ± SD: 21.5 ± 1.5 years; 18 female and 46 male). Handedness was evaluated using the Edinburgh Handedness Inventory^71^. The participants had normal or corrected-to-normal vision, normal hearing abilities, and reported no history of neurological or psychiatric conditions. Thirty-two pairs of participants of the same sex who did not know each other before the experiment were formed.

This study was approved by the Ethics Committee of Tohoku University Graduate School of Medicine, and was conducted according to the Declaration of Helsinki. Written informed consent was obtained from all participants. Participants received monetary compensation for their participation.

### Experimental procedure

Each experimental session consisted of four steps (Fig. 1a). During the first two steps in a session, a pair of participants was seated facing each other and underwent (1) a *rhythmic movement block* in which they moved their right arms in sync to a ticking sound presented respectively at a constant tempo, and (2) a *word teaching-learning block* in which they took turns teaching and learning given unknown English words. The participants were then separated and answered (3) a questionnaire regarding the subjective rapport they felt during the teaching-learning block, and (4) a paper test on the taught/learnt words during that session. Details of each step are provided below.

Each pair performed two sessions, one with a *synchronous condition* and one with an *asynchronous condition* in the rhythmic movement block. The order of the conditions for the two sessions was counterbalanced across the participant pairs. Between the two sessions, participants took a 5-minute rest or more if either of the pair requested it.

In order to probe into the effect of *incidental* physical synchrony, before the experiment, we provided participants with a story claiming that the purpose of the study was “to investigate the effect of embodied physical tempo on the succeeding learning process”. In the debriefing, after all procedures were completed, the true aim of the study was explained and was confirmed to have been unknown to all participants.

#### Rhythmic movement block

During the first step of an experimental session, each participant pair performed a repetitive arm movement task of 6 minutes, sitting on opposite sides of a table, facing each other. A two-beat sound (alternating high- and low-pitch beeps) with a constant tempo of 23, 26, or 29 beats per minutes (BPM) was presented to the participants through earphones. Participants were instructed to put and keep their right elbow on a cushioned block on the table, and to tilt the right forearm inwards (i.e. to the left) at the timing of each high-pitch beat, then move it back out at the timing of each low-pitch beat. In the synchronous (“Sync”) condition, the beat sound presented to the pair had the same tempo and no phase difference, so that the pair was conducting the movement in sync. In the asynchronous (“Async”) condition, the tempo of the beat sound was different for the two participants. They were instructed to focus on the beat sound and move their arms precisely following the timing. They were also instructed to keep their eyes fixated on the tip of a pole (height 37 cm from the table top) that was put in the middle between them, with the plausible purpose of “preventing unwanted head movements”. By this setting, each other’s arm movements were within their eyesight.

The tempi were counterbalanced across participant pairs. Within each pair, the tempo for the synchronous condition and the two tempi for the asynchronous condition were different. Supplementary Table S1 presents further information on the assignment of conditions and tempi.

#### Word teaching-learning block

After the rhythmic movement block, the earphones were removed, and a pile of eight “scenario cards” for the teaching-learning task was placed at each participant’s right-hand side on the table. The following word teaching-learning block took 6 minutes and 10 seconds.

A block contained eight trials of 40-second duration, with 5 seconds of inter-trial intervals (ITIs) between trials. The beginning and ending of the trials were cued by beep sounds from a programmed timer. During a trial, one of the pair was assigned as the teacher and the other as the student. At the cue, they picked a scenario card from their respective pile with their right hand, and held it next to their right eye (not in front, in order to avoid blocking their partner’s sight). The teacher’s card contained an English word, its pronunciation, its meaning in Japanese, and an example sentence in English. The teacher repeatedly delivered the word, meaning, and example sentence to the student, while trying to memorize everything for themselves. No ad-libs were allowed. The student’s card contained only the word and its pronunciation. The student silently listened to the teacher and tried to memorize the information, while confirming the word on the card if needed. No questions or comments were allowed. Their respective role (teacher or student) was also indicated on their cards. At the cue, they put the scenario cards on the table face down, and took a break during the 5-second ITI. There were also 5-second breaks at the beginning and end of the block.

At the beginning of the experiment, participants were explained that the goal of the teaching-learning task was to maximize one’s own and the partner’s total score in the following word memory test (see below), and were instructed to try their best so that both of them could obtain the highest possible scores.

The words were selected from the GRE (graduate record examination) word list and were expected to be unknown to most participants. We prepared two sets of eight words, one starting with the letter *a* and the other with the letter *c*. The complete list of the used words and the example sentences are shown in Supplementary Table S2.

#### Rapport and learning questionnaire

After the end of each session, a pair was separated, seated at different desks, and completed a questionnaire on their subjective rapport and impression of the learning process during the teaching-learning block. The questionnaire consisted of nine items: (1) I was able to explain to the partner in an easy-to-understand way; (2) “The partner’s explanation was easy to understand”; (3) “I was focusing my attention to the partner”; (4) “I felt the partner was focusing his/her attention to me”; (5) “I had a favourable impression on the partner”; (6) “I felt the partner was friendly to me”; (7) “I felt that my actions and the partner’s actions were in harmony”; (8) “I felt united with the partner”; and (9) “I feel confident on the outcome of the teaching-learning task”. Six items measured the three-component model of rapport^56^: mutual attentiveness (items 3, 4); positivity (items 5, 6); and coordination (items 7, 8). These items were identified as representative of the three components in the previously developed questionnaires on rapport in face-to-face interactions^13, 72–74^. The additional three items (1, 2, and 9) probed into the perceptions of one’s own and the partner’s teaching process and into the participants’ confidence in the learning outcome. Participants were instructed to rate the teaching-learning interaction regarding the nine items, on a seven-level Likert scale from 1 = “strongly disagree” to 7 = “strongly agree”. No time limit was imposed for the questionnaire, but the participants generally completed it within a few minutes.

#### Word memory test

After both participants in a pair completed the rapport questionnaire, they separately took a paper test on the eight taught/learnt words in that session. On the test sheet, the eight words were presented in a shuffled order (i.e. different from the presented order in the teaching-learning block), and the participants were asked to fill in the meaning in Japanese and the example sentence in English for each word. For the example sentences, they were instructed that it was not necessary to reproduce the exact sentence that was taught/learnt during the teaching-learning block but that scores would only be given to the answer sentences that indicated comprehension of the correct meaning. They were also requested to check whether each word was “known”, “seen somewhere before but not remembered”, or “unknown” to them. They were notified when 5 minutes had passed, and the answer paper was collected at the time limit of 6 minutes. Then the pair took a rest for 5 minutes, or longer if they requested. They were explicitly told that the words in the finished sessions would not reappear in the following sessions and that there was thus no need to try to remember them.

#### Practice session

After receiving explanations on the procedure and before starting the first real session, each pair conducted a short practice session, which consisted of (1) a 3-minute rhythmic movement block with an asynchronous condition using 23 BPM and 29 BPM (fixed for all pairs), (2) a 3-minute word teaching-learning block in which four instead of eight words were taught/learnt, (3) the rapport questionnaire which was same as the one used in the real sessions, and (4) a word memory test on the four words with a shortened time limit of 3 minutes. We checked that participants understood and conformed to the procedures in each block.

### fNIRS measurements

Prefrontal neural activities during the word teaching-learning blocks were measured using HOT-1000 (Hitachi High-Technologies, Japan), a wireless continuous-wave fNIRS device (Fig. 1b). This device has two horizontally placed sets of dual source-detector (SD) optode pairs, each consisting of one light source and two light detectors that are placed at the distance of 1 cm and 3 cm from the light source. Similarly to the device used in previous studies^26, 50^, the light source of the fNIRS device uses only a single wavelength of 810 nm and the concentration change of total haemoglobin (totalHb) on the optical path is estimated for each of the two SD pairs with distances of 1.0 cm and 3.0 cm. The 1-cm-SD pair provides an auxiliary measurement signal from shallow tissues like scalp and skull, and is not affected by the blood flow changes in the underlying cortex^58, 59^. This signal is called a “shallow signal”, and was used to extract the brain signal from the “deep signal” from the 3-cm-SD channel and to confirm the neural origin of the observed IBS (see below). Gagnon *et al.* suggested that totalHb changes are less affected by pial vein contamination and provide better spatial specificity than oxygenated or deoxygenated haemoglobin changes^75^. A considerable number of fNIRS-hyperscanning studies have also used the totalHb changes^76, 48, 26, 50^, supporting that our analyses based on totalHb measurements are sufficiently sensitive. The horizontal positions of the two optode sets are adjustable. We placed them at the right medial (with the 3-cm detector on the Fpz position defined by the international 10-20 system for electroencephalography electrode placement) and the left lateral prefrontal regions (with the 3-cm detector 3 cm left from the Fpz; see Fig. 1b). Based on the virtual registration method^77, 78^, the right 3-cm-SD channel corresponded to the medial part of the Brodmann area (BA)10, and the left 3-cm-SD channel to the border of the dorsolateral prefrontal cortex (BA46) and the lateral part of BA10. We chose these regions with the intention to assess more externally- and internally-oriented neural functions by the lateral and medial probes, respectively. The participants’ hair under the optodes was carefully pushed off their foreheads, and the optodes were covered with a rubber cover to block out light.

Although our focus was on IBS during the word teaching-learning block, to make the story about the study purpose (“investigating the effect of embodied tempo on learning”) and the instruction to keep their fixation during the rhythmic movement block more credible for the participants, we measured brain activities throughout both blocks. In order to minimize motion noises, participants were instructed to adopt a relaxed posture in the seat and to maintain it, and to avoid head movements as far as possible. Participants were especially instructed to only move their eyes instead of looking down and up during the teaching-learning block.

### Data analysis

#### Evaluation of rapport and memory scores

Rapport scores for each dyad were calculated by averaging the ratings for the six questionnaire items (items 3-8) of the two members. For the post-hoc analyses of the three components of the rapport, ratings of the corresponding two items (see above) were averaged within each pair, respectively. The scores thus ranged from 1 to 7.

For scoring the word memory test, the words classified as “known” and “have seen” were excluded, and the accuracy for the remaining “unknown” words was calculated. Only 2.3% of the words were excluded at the ground level, and only one word at maximum was excluded at the individual level. For each word, 0.5 points were given if the meaning was correct, and 0 points otherwise. Another 0.5 points were given if the example sentence was correct, 0 points otherwise. Grammatical and spelling errors were not taken into account. For each session, the score was averaged within each individual, and then averaged within each pair. The scores thus ranged from 0 to 1.

As a supplementary analysis, as a measure of subjectively perceived effectiveness of learning, the ratings for the questionnaire items 1, 2, 9 were averaged within each pair, and analysed in the same manner as the word memory test scores. The scores ranged from 1 to 7.

In order to cancel the potential influence of familiarization over the sessions and the difference in difficulty of the word sets taught/learnt, effects of session order were adjusted by subtracting the between-session differences in group averages for these teaching-learning communication outcomes (note that the word sets were fixed to sessions, so there was no need to adjust their effects separately).

#### fNIRS data preprocessing

The fNIRS signals were first detrended and low-pass filtered (cut-off at 0.2 Hz) to reduce potential instrumental drift and high-frequency physiological noises^79, 80^. Then, to reduce the influence of spike-like artefacts induced by head motion or other causes, a wavelet-based motion artefact reduction method^81, 82^ implemented in HomER2^83^ (http://www.nmr.mgh.harvard.edu/PMI/resources/homer2/home.htm) was applied. Finally, dual SD regression was applied to dissociate brain-originated signal from systemic and motion-related noises by regressing out the shallow signal component provided by the 1-cm-SD channels from the deep signals from 3-cm-SD channels^84–87^.

#### Evaluation of IBS

To evaluate the coherence of the fNIRS signals between the pairs of participants, we used wavelet transform coherence (WTC), a method to evaluate a localized correlation coefficient in time-frequency space^57^. A cross wavelet and wavelet coherence toolbox for MATLAB (http://noc.ac.uk/using-science/crosswavelet-wavelet-coherence) was used for calculation. The IBS values were calculated by averaging the WTC *R*^2^ values at the Fourier period corresponding to the task cycle of 45 seconds (= 40 seconds of teaching-learning communication and 5 seconds of ITI) over the whole teaching-learning block, except for the time points within the cone of influence to avoid contaminating influences from edge effects^88^.

In the same way as for the rapport and word memory scores, the potential influence of familiarization over the sessions and the differences in the word sets were cancelled by adjusting the session order effects.

#### Statistical tests

The hypothesized enhancing effects of prior physical synchrony in comparison to asynchrony on the communication outcomes (hypothesis 1) and IBS (hypothesis 2) were tested using paired t-tests. For testing the positive association between the rapport and left lateral PFC IBS that both showed enhancement induced by prior physical synchrony (hypothesis 3), taking their hierarchical nature due to the within-subject design into account, the repeated measure correlation^89^ was used. This analysis captures within-unit relationships between two variables, and, for the test of association, is equivalent to the linear mixed-effects model with centering within-cluster for the independent variable.

To correct for multiple testing, false discovery rate (FDR) control via the adaptive Benjamini-Hochberg procedure^90^ was used. P-values were FDR-adjusted for the tests of the main hypotheses 1-3. P-values for the post-hoc analyses on the three components rapport were separately FDR-adjusted. P-values for the t-tests on the shallow signals and on the signals without correcting for shallow components were also adjusted separately, because these tests were conducted to confirm that the observed IBS enhancement was not due to the artefacts reflected in the shallow signals (i.e. opposite to the main hypotheses, the null hypothesis was not expected to be rejected). Results with FDR-adjusted *p*_FDR_ < 0.05 were deemed significant.

## Acknowledgements

We thank Ms. Y. Yamada for helping with our experiment, the participants in this study, and all colleagues in our laboratory for their support. This work was partially supported by JSPS KAKENHI Grant Numbers JP26330171, JP15H01771, and JP17H01753, and also by the Center of Innovation Program from the Japan Science and Technology Agency (JST), Japan.

## References

1. Marsh, K. L., Richardson, M. J. & Schmidt, R. Social connection through joint action and interpersonal coordination. Topics in Cognitive Science 1, 320–339 (2009).

2. Chartrand, T. L. & Bargh, J. A. The chameleon effect: the perception–behavior link and social interaction. J Pers Soc Psychol 76, 893 (1999).

3. Rennung, M. & Göritz, A. S. Prosocial consequences of interpersonal synchrony: A meta-analysis. Zeitschrift fur Psychologie 224, 168–189 (2016).

4. Vicaria, I. M. & Dickens, L. Meta-analyses of the intra- and interpersonal outcomes of interpersonal coordination. Journal of Nonverbal Behavior 40, 335–361 (2016).

5. Hove, M. J. & Risen, J. L. It’s all in the timing: Interpersonal synchrony increases affiliation. Social Cognition 27, 949–960 (2009).

6. Wiltermuth, S. S. & Heath, C. Synchrony and cooperation. Psychological Science 20, 1–5 (2009).

7. Valdesolo, P. & DeSteno, D. Synchrony and the social tuning of compassion. Emotion 11, 262 (2011).

8. Valdesolo, P., Ouyang, J. & DeSteno, D. The rhythm of joint action: Synchrony promotes cooperative ability. J Exp Soc Psychol 46, 693–695 (2010).

9. Tschacher, W., Rees, G. M. & Ramseyer, F. Nonverbal synchrony and affect in dyadic interactions. Front Psychol 5, 1323 (2014).

10. Macrae, C. N., Duffy, O. K., Miles, L. K. & Lawrence, J. A case of hand waving: Action synchrony and person perception. Cognition 109, 152–156 (2008).

11. Bernieri, F. J. Coordinated movement and rapport in teacher-student interactions. Journal of Nonverbal behavior 12, 120–138 (1988).

12. LaFrance, M. & Broadbent, M. Group rapport: Posture sharing as a nonverbal indicator. Group & Organization Studies 1, 328–333 (1976).

13. Zhou, J. The effects of reciprocal imitation on teacher–student relationships and student learning outcomes. Mind, Brain, and Education 6, 66–73 (2012).

14. Rabinowitch, T.-C. & Meltzoff, A. N. Synchronized movement experience enhances peer cooperation in preschool children. J Exp Child Psychol 160, 21–32 (2017).

15. Ramseyer, F. & Tschacher, W. Nonverbal synchrony of head- and body-movement in psychotherapy: different signals have different associations with outcome. Front Psychol 5, 979 (2014).

16. Velandia, R. The role of warming up activities in adolescent students’ involvement during the English class. Profile 10, 9–26 (2008). Profile Issues in Teachers Professional Development.

17. Kent, A. Synchronization as a classroom dynamic: A practitioner’s perspective. Mind, Brain, and Education 7, 13–18 (2013).

18. Engh, D. Why use music in English language learning? a survey of the literature. English Language Teaching 6, 113 (2013).

19. Konvalinka, I. & Roepstorff, A. The two-brain approach: how can mutually interacting brains teach us something about social interaction? Front Hum Neurosci 6, 215 (2012).

20. Schilbach, L. et al. Toward a second-person neuroscience. Behav Brain Sci 36, 393–414 (2013).

21. Scholkmann, F., Holper, L., Wolf, U. & Wolf, M. A new methodical approach in neuroscience: assessing inter-personal brain coupling using functional near-infrared imaging (fNIRI) hyperscanning. Front Hum Neurosci 7, 813 (2013).

22. Babiloni, F. & Astolfi, L. Social neuroscience and hyperscanning techniques: past, present and future. Neurosci Biobehav Rev 44, 76–93 (2014).

23. Koike, T., Tanabe, H. C. & Sadato, N. Hyperscanning neuroimaging technique to reveal the “two-in-one” system in social interactions. Neurosci Res 90, 25–32 (2015).

24. Hari, R., Himberg, T., Nummenmaa, L., Hämäläinen, M. & Parkkonen, L. Synchrony of brains and bodies during implicit interpersonal interaction. Trends Cogn Sci 17, 105–106 (2013).

25. Cui, X., Bryant, D. M. & Reiss, A. L. NIRS-based hyperscanning reveals increased interpersonal coherence in superior frontal cortex during cooperation. Neuroimage 59, 2430–2437 (2012).

26. Nozawa, T., Sasaki, Y., Sakaki, K., Yokoyama, R. & Kawashima, R. Interpersonal frontopolar neural synchronization in group communication: An exploration toward fNIRS hyperscanning of natural interactions. Neuroimage 133, 484–497 (2016).

27. Liu, N. et al. NIRS-based hyperscanning reveals inter-brain neural synchronization during cooperative jenga game with face-to-face communication. Front Hum Neurosci 10, 82 (2016).

28. Hu, Y. et al. Inter-brain synchrony and cooperation context in interactive decision making. Biol Psychol 133, 54–62 (2018).

29. Miller, J. G. et al. Inter-brain synchrony in mother-child dyads during cooperation: An fnirs hyperscanning study. Neuropsychologia 124, 117–124 (2019).

30. Jiang, J. et al. Neural synchronization during face-to-face communication. J Neurosci 32, 16064–16069 (2012).

31. Tang, H. et al. Interpersonal brain synchronization in the right temporo-parietal junction during face-to-face economic exchange. Soc Cogn Affect Neurosci 11, 23–32 (2016).

32. Schippers, M. B., Roebroeck, A., Renken, R., Nanetti, L. & Keysers, C. Mapping the information flow from one brain to another during gestural communication. Proc Natl Acad Sci U S A 107, 9388–9393 (2010).

33. Stephens, G. J., Silbert, L. J. & Hasson, U. Speaker-listener neural coupling underlies successful communication. Proc Natl Acad Sci U S A 107, 14425–14430 (2010).

34. Cheng, X., Li, X. & Hu, Y. Synchronous brain activity during cooperative exchange depends on gender of partner: A fNIRS-based hyperscanning study. Hum Brain Mapp 36, 2039–2048 (2015).

35. Bilek, E. et al. Information flow between interacting human brains: Identification, validation, and relationship to social expertise. Proc Natl Acad Sci U S A 112, 5207–5212 (2015).

36. Liu, Y. et al. Measuring speaker-listener neural coupling with functional near infrared spectroscopy. Sci Rep 7, 43293 (2017).

37. Anders, S., Heinzle, J., Weiskopf, N., Ethofer, T. & Haynes, J.-D. Flow of affective information between communicating brains. Neuroimage 54, 439–446 (2011).

38. Koike, T. et al. Neural substrates of shared attention as social memory: A hyperscanning functional magnetic resonance imaging study. Neuroimage 125, 401–412 (2016).

39. Pan, Y., Cheng, X., Zhang, Z., Li, X. & Hu, Y. Cooperation in lovers: An fNIRS-based hyperscanning study. Hum Brain Mapp 38, 831–841 (2017).

40. Koide, T. & Shimada, S. Cheering enhances inter-brain synchronization between sensorimotor areas of player and observer. Japanese Psychological Research 60, 265–275 (2018).

41. Watanabe, K. Teaching as a dynamic phenomenon with interpersonal interactions. Mind, Brain, and Education 7, 91–100 (2013).

42. Holper, L. et al. The teaching and the learning brain: A cortical hemodynamic marker of teacher–student interactions in the socratic dialog. International Journal of Educational Research 59, 1–10 (2013).

43. Takeuchi, N., Mori, T., Suzukamo, Y. & Izumi, S.-I. Integration of teaching processes and learning assessment in the prefrontal cortex during a video game teaching–learning task. Front Psychol 7, 2052 (2017).

44. Dikker, S. et al. Brain-to-brain synchrony tracks real-world dynamic group interactions in the classroom. Curr Biol 27, 1375–1380 (2017).

45. Bevilacqua, D. et al. Brain-to-brain synchrony and learning outcomes vary by student–teacher dynamics: Evidence from a real-world classroom electroencephalography study. J Cognit Neurosci 1–11 (2018).

46. Zheng, L. et al. Enhancement of teaching outcome through neural prediction of the students’ knowledge state. Hum Brain Mapp 39, 3046–3057 (2018).

47. Yun, K., Watanabe, K. & Shimojo, S. Interpersonal body and neural synchronization as a marker of implicit social interaction. Sci Rep 2, 959 (2012).

48. Holper, L., Scholkmann, F. & Wolf, M. Between-brain connectivity during imitation measured by fNIRS. Neuroimage 63, 212–222 (2012).

49. Osaka, N. et al. How two brains make one synchronized mind in the inferior frontal cortex: fNIRS-based hyperscanning during cooperative singing. Front Psychol 6, 1811 (2015).

50. Ikeda, S. et al. Steady beat sound facilitates both coordinated group walking and inter-subject neural synchrony. Front Hum Neurosci 11, 147 (2017).

51. Amodio, D. M. & Frith, C. D. Meeting of minds: the medial frontal cortex and social cognition. Nat Rev Neurosci 7, 268–277 (2006).

52. Gilbert, S. J. et al. Functional specialization within rostral prefrontal cortex (area 10): a meta-analysis. J Cogn Neurosci 18, 932–948 (2006).

53. Bludau, S. et al. Cytoarchitecture, probability maps and functions of the human frontal pole. Neuroimage 93, 260–275 (2014).

54. Ferrari, M. & Quaresima, V. A brief review on the history of human functional near-infrared spectroscopy (fNIRS) development and fields of application. Neuroimage 63, 921–935 (2012).

55. Scholkmann, F. et al. A review on continuous wave functional near-infrared spectroscopy and imaging instrumentation and methodology. Neuroimage 85 Pt 1, 6–27 (2014).

56. Tickle-Degnen, L. & Rosenthal, R. The nature of rapport and its nonverbal correlates. Psychological inquiry 1, 285–293 (1990).

57. Grinsted, A., Moore, J. C. & Jevrejeva, S. Application of the cross wavelet transform and wavelet coherence to geophysical time series. Nonlinear Processes in Geophysics 11, 561–566 (2004).

58. Fukui, Y., Ajichi, Y. & Okada, E. Monte Carlo prediction of near-infrared light propagation in realistic adult and neonatal head models. Appl Opt 42, 2881–2887 (2003).

59. Strangman, G. E., Zhang, Q. & Li, Z. Scalp and skull influence on near infrared photon propagation in the colin27 brain template. Neuroimage 85 Pt 1, 136–149 (2014).

60. Cacioppo, S. et al. You are in sync with me: neural correlates of interpersonal synchrony with a partner. Neuroscience 277, 842–858 (2014).

61. Neubert, F.-X., Mars, R. B., Thomas, A. G., Sallet, J. & Rushworth, M. F. S. Comparison of human ventral frontal cortex areas for cognitive control and language with areas in monkey frontal cortex. Neuron 81, 700–713 (2014).

62. Moayedi, M., Salomons, T. V., Dunlop, K. A., Downar, J. & Davis, K. D. Connectivity-based parcellation of the human frontal polar cortex. Brain Struct Funct 220, 2603–2616 (2015).

63. Gilbert, S. J., Gonen-Yaacovi, G., Benoit, R. G., Volle, E. & Burgess, P. W. Distinct functional connectivity associated with lateral versus medial rostral prefrontal cortex: a meta-analysis. Neuroimage 53, 1359–1367 (2010).

64. Peng, K., Steele, S. C., Becerra, L. & Borsook, D. Brodmann area 10: Collating, integrating and high level processing of nociception and pain. Prog Neurobiol 161, 1–22 (2018).

65. Koechlin, E. & Hyafil, A. Anterior prefrontal function and the limits of human decision-making. Science 318, 594–598 (2007).

66. Kouneiher, F., Charron, S. & Koechlin, E. Motivation and cognitive control in the human prefrontal cortex. Nat Neurosci 12, 939–945 (2009).

67. Van Overwalle, F. Social cognition and the brain: a meta-analysis. Hum Brain Mapp 30, 829–858 (2009).

68. Clark, I. & Dumas, G. Toward a neural basis for peer-interaction: what makes peer-learning tick? Front Psychol 6, 28 (2015).

69. Jiang, J. et al. Leader emergence through interpersonal neural synchronization. Proc Natl Acad Sci U S A 112, 4274–4279 (2015).

70. Kinreich, S., Djalovski, A., Kraus, L., Louzoun, Y. & Feldman, R. Brain-to-brain synchrony during naturalistic social interactions. Sci Rep 7, 17060 (2017).

71. Oldfield, R. C. The assessment and analysis of handedness: the Edinburgh inventory. Neuropsychologia 9, 97–113 (1971).

72. Bernieri, F. J., Gillis, J. S., Davis, J. M. & Grahe, J. E. Dyad rapport and the accuracy of its judgment across situations: a lens model analysis. Journal of Personality and Social Psychology 71, 110 (1996).

73. Puccinelli, N. M. & Tickle-Degnen, L. Knowing too much about others: Moderators of the relationship between eavesdropping and rapport in social interaction. Journal of Nonverbal Behavior 28, 223–243 (2004).

74. Lakens, D. & Stel, M. If they move in sync, they must feel in sync: Movement synchrony leads to attributions of rapport and entitativity. Social Cognition 29, 1–14 (2011).

75. Gagnon, L. et al. Quantification of the cortical contribution to the NIRS signal over the motor cortex using concurrent NIRS-fMRI measurements. Neuroimage 59, 3933–3940 (2012).

76. Dommer, L., Jäger, N., Scholkmann, F., Wolf, M. & Holper, L. Between-brain coherence during joint n-back task performance: a two-person functional near-infrared spectroscopy study. Behav Brain Res 234, 212–222 (2012).

77. Tsuzuki, D. et al. Virtual spatial registration of stand-alone fNIRS data to MNI space. Neuroimage 34, 1506–1518 (2007).

78. Okamoto, M. et al. Three-dimensional probabilistic anatomical cranio-cerebral correlation via the international 10-20 system oriented for transcranial functional brain mapping. Neuroimage 21, 99–111 (2004).

79. Katura, T., Tanaka, N., Obata, A., Sato, H. & Maki, A. Quantitative evaluation of interrelations between spontaneous low-frequency oscillations in cerebral hemodynamics and systemic cardiovascular dynamics. Neuroimage 31, 1592–1600 (2006).

80. Matthews, F., Pearlmutter, B. A., Ward, T. E., Soraghan, C. & Markham, C. Hemodynamics for brain-computer interfaces: optical correlates of control signals. IEEE Signal Processing Magazine 25, 87–94 (2008).

81. Molavi, B. & Dumont, G. A. Wavelet-based motion artifact removal for functional near-infrared spectroscopy. Physiol Meas 33, 259–270 (2012).

82. Brigadoi, S. et al. Motion artifacts in functional near-infrared spectroscopy: a comparison of motion correction techniques applied to real cognitive data. Neuroimage 85 Pt 1, 181–191 (2014).

83. Huppert, T. J., Diamond, S. G., Franceschini, M. A. & Boas, D. A. HomER: a review of time-series analysis methods for near-infrared spectroscopy of the brain. Applied Optics 48, D280–D298 (2009).

84. Toronov, V. et al. Study of local cerebral hemodynamics by frequency-domain near-infrared spectroscopy and correlation with simultaneously acquired functional magnetic resonance imaging. Opt Express 9, 417–427 (2001).

85. Robertson, F., Douglas, T. & Meintjes, E. Motion artefact removal for functional near infrared spectroscopy: a comparison of methods. IEEE Trans Biomed Eng 57, 1377–1387 (2010).

86. Saager, R. B., Telleri, N. L. & Berger, A. J. Two-detector corrected near infrared spectroscopy (C-NIRS) detects hemodynamic activation responses more robustly than single-detector NIRS. Neuroimage 55, 1679–1685 (2011).

87. Scholkmann, F., Metz, A. J. & Wolf, M. Measuring tissue hemodynamics and oxygenation by continuous-wave functional near-infrared spectroscopy—how robust are the different calculation methods against movement artifacts? Physiol Meas 35, 717 (2014).

88. Torrence, C. & Compo, G. P. A practical guide to wavelet analysis. Bulletin of the American Meteorological society 79, 61–78 (1998).

89. Bakdash, J. Z. & Marusich, L. R. Repeated measures correlation. Front Psychol 8, 456 (2017).

90. Benjamini, Y. & Hochberg, Y. On the adaptive control of the false discovery rate in multiple testing with independent statistics. Journal of educational and Behavioral Statistics 25, 60–83 (2000).

91. Xia, M., Wang, J. & He, Y. BrainNet Viewer: a network visualization tool for human brain connectomics. PLoS One 8, e68910 (2013).

