## Supplementary Information for "Prior physical synchrony enhances rapport and inter-brain synchronization during subsequent educational communication"

**Manuscript title:**

### Supplementary Tables

**Table S1. Assignment of physical synchronous/asynchronous conditions and the pairs of tempi for the rhythmic movement block of each session for each participant pair.**

| Pair No. | Session 1 | | Session 2 | |
|---|---|---|---|---|
| | Condition | Tempo pair (BPM) | Condition | Tempo pair (BPM) |
| 1 | Sync | (23, 23) | Async | (26, 29) |
| 2 | Async | (29, 26) | Sync | (23, 23) |
| 3 | Sync | (26, 26) | Async | (23, 29) |
| 4 | Async | (29, 23) | Sync | (26, 26) |
| 5 | Async | (26, 29) | Sync | (23, 23) |
| 6 | Sync | (29, 29) | Async | (23, 26) |
| 7 | Sync | (23, 23) | Async | (29, 26) |
| 8 | Async | (26, 23) | Sync | (29, 29) |
| 9 | Async | (23, 29) | Sync | (26, 26) |
| 10 | Sync | (26, 26) | Async | (29, 23) |
| 11 | Sync | (23, 23) | Async | (26, 29) |
| 12 | Async | (23, 26) | Sync | (29, 29) |
| 13 | Async | (29, 26) | Sync | (23, 23) |
| 14 | Sync | (29, 29) | Async | (26, 23) |
| 15 | Sync | (26, 26) | Async | (23, 29) |
| 16 | Async | (29, 23) | Sync | (26, 26) |
| 17 | Async | (26, 29) | Sync | (23, 23) |
| 18 | Sync | (29, 29) | Async | (23, 26) |
| 19 | Sync | (23, 23) | Async | (29, 26) |
| 20 | Async | (26, 23) | Sync | (29, 29) |
| 21 | Async | (23, 29) | Sync | (26, 26) |
| 22 | Sync | (26, 26) | Async | (29, 23) |
| 23 | Async | (23, 26) | Sync | (29, 29) |
| 24 | Sync | (23, 23) | Async | (26, 29) |
| 25 | Sync | (29, 29) | Async | (26, 23) |
| 26 | Async | (29, 26) | Sync | (23, 23) |
| 27 | Sync | (26, 26) | Async | (23, 29) |
| 28 | Async | (29, 23) | Sync | (26, 26) |
| 29 | Sync | (23, 23) | Async | (29, 26) |
| 30 | Async | (26, 23) | Sync | (29, 29) |
| 31 | Async | (23, 29) | Sync | (26, 26) |
| 32 | Sync | (29, 29) | Async | (26, 23) |

**Table S2. Words and example sentences used in the word teaching-learning blocks of the two sessions.**

*For Session 1*

| Word | Example sentence |
|---|---|
| abrogate | The country may abrogate the international agreement. |
| abstain | It is often difficult to abstain from drinking. |
| acrid | The room was filled with the acrid smell of tobacco. |
| adroit | He is adroit in making excuses. |
| ameliorate | Judge ordered the company to ameliorate working conditions. |
| appease | Only a sincere apology will appease my anger. |
| astute | The boss made an astute business decision. |
| austere | The church was austere and simple. |

*For Session 2*

| Word | Example sentence |
|---|---|
| cajole | The salesman will cajole you into buying the car. |
| callow | He is a young callow man with no experience. |
| candid | He is trusted because he always gives candid opinions. |
| capitulate | He capitulated to his wife's demand. |
| caustic | The teacher was known by his caustic comments. |
| censure | The food factory was censured for its carelessness. |
| choleric | He is friendly at one moment, choleric the next. |
| cringe | Large noise of thunder made the children cringe. |

**Supplementary Figures**

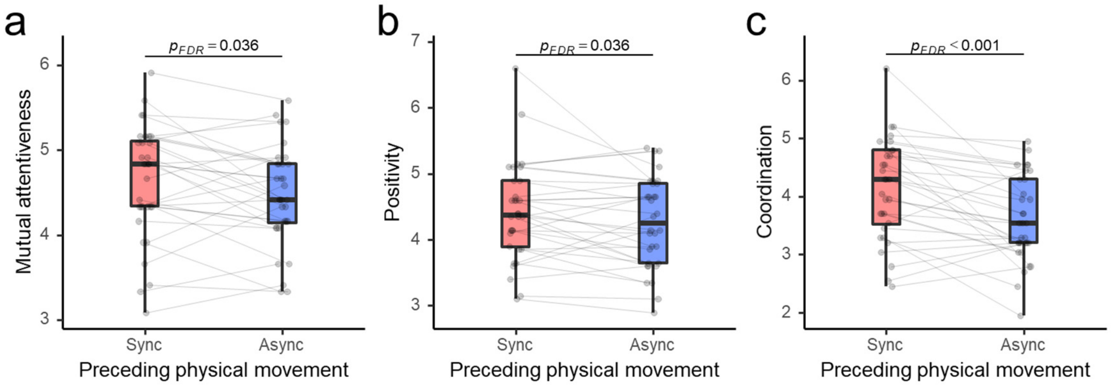


**Figure S1. Three components of rapport, (a) mutual attentiveness, (b) positivity, and (c) coordination, during the word teaching-learning task after experiencing physical synchrony/asynchrony.**

Box plots show the median, interquartile range (IQR), and minimum/maximum values of the average rapport ratings during the teaching-learning block of all dyads for the two conditions of the preceding rhythmic movement block. Grey points with connecting lines represent the two conditions of each dyad.

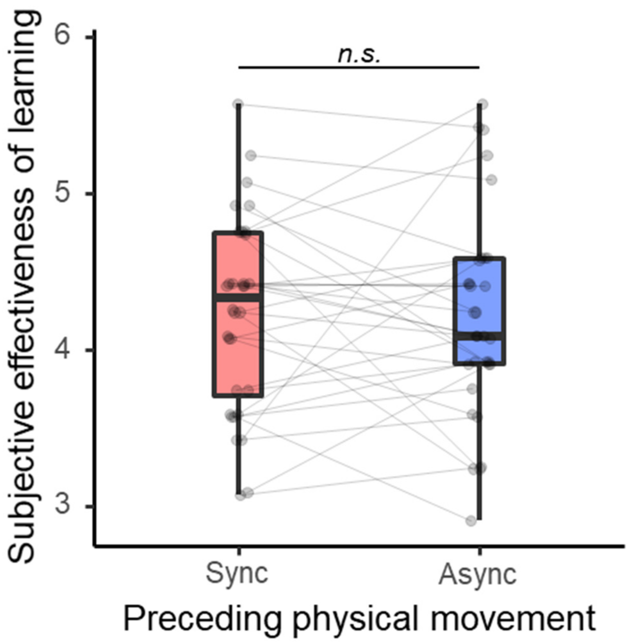


**Figure S2. Subjective effectiveness of learning after experiencing physical synchrony/asynchrony.**

See the caption of Fig. S1 for box plot notations.

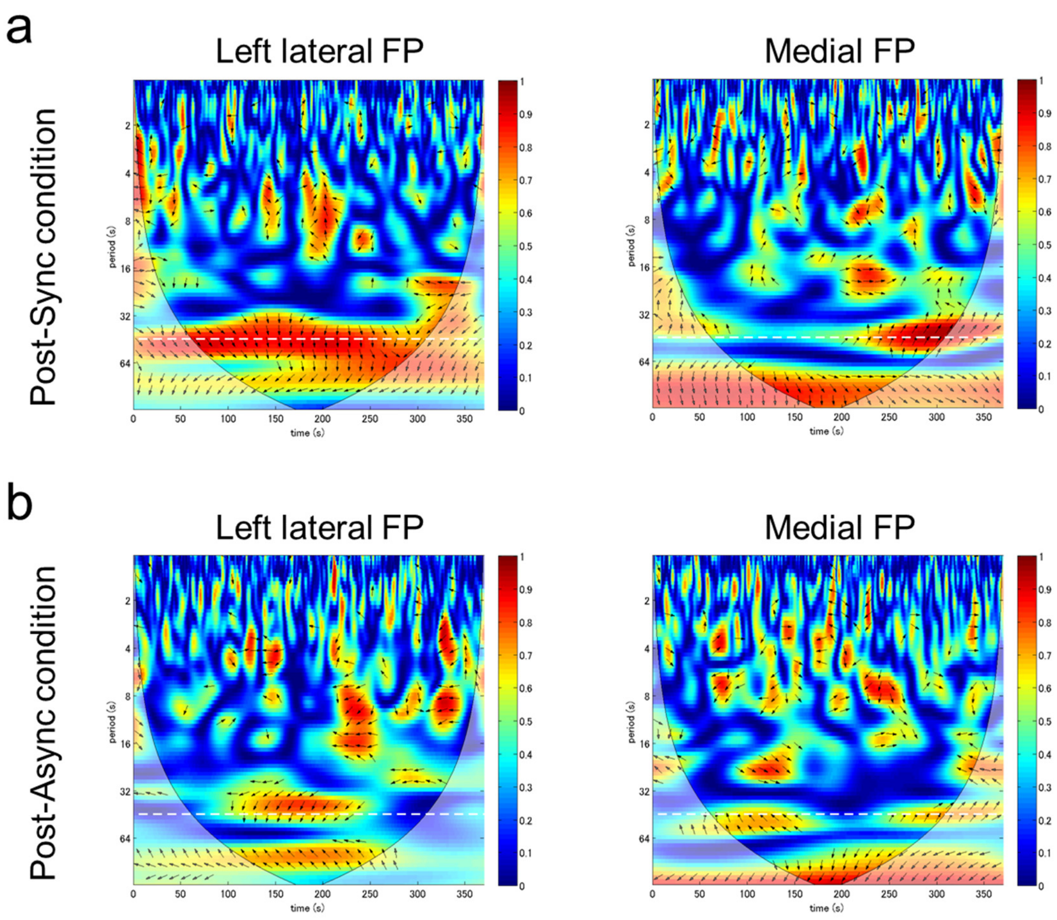


**Figure S3. Illustrative examples of wavelet transform coherence (WTC) of neural signals from a single pair of participants.**

Left and right panels show interpersonal coherence of the left lateral and medial PFC, respectively, during the word teaching-learning task after experiencing physical synchrony (a) and asynchrony (b). In each panel, the horizontal white dashed line at period = 45 seconds indicates the teaching-learning task period, at which WTC values were analysed. The faded colour areas bordered by spindle-like curves indicate the cones of influence (COIs), where the values can suffer from edge effects and thus were excluded from the analysis (see main text). In the areas with high coherence values, arrows indicate phase angle of the paired signals, with right and left directions correspond to in-phase (0) and anti-phase (π), respectively.

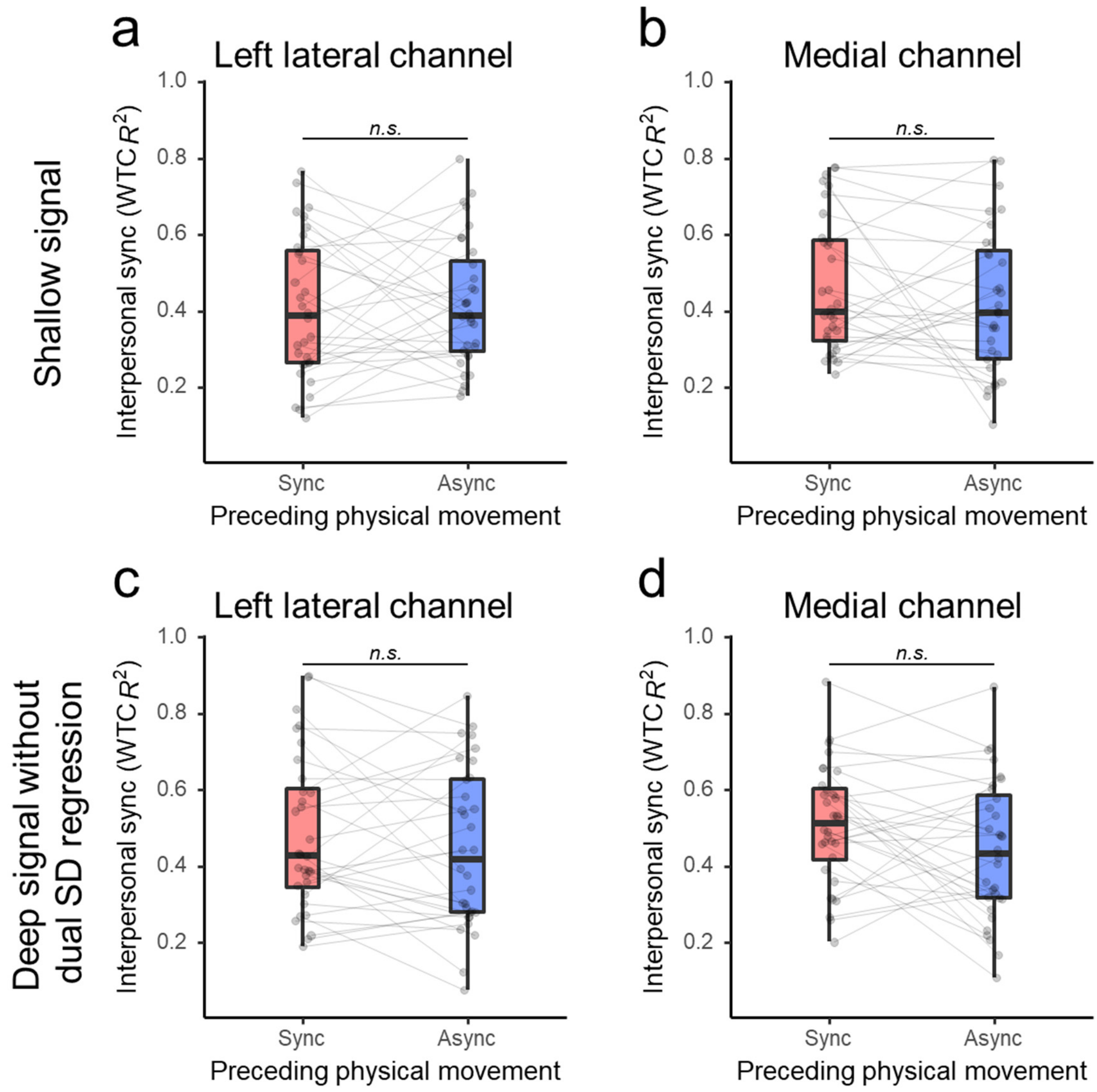


**Figure S4. Interpersonal synchronization values of contaminating and contaminated signals.**

Values were calculated for the shallow signals from the left lateral (a) and medial (b) 1-cm-SD channels, and for the deep signals from the left lateral (c) and medial (d) 3-cm-SD channels without the removal of shallow signal components by dual SD regression, during the word teaching-learning task after experiencing physical synchrony/asynchrony. See the caption of Fig. S1 for box plot notations.

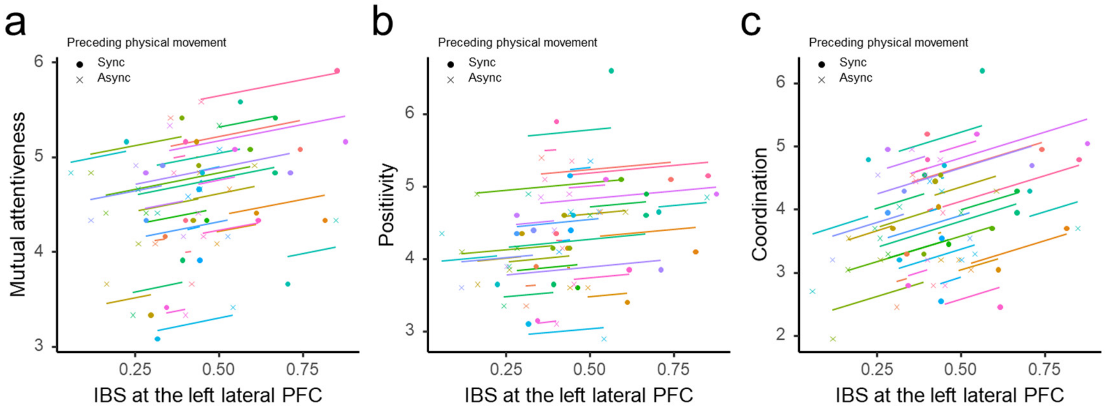


**Figure S5. Within-pair correlation between IBS at the left lateral prefrontal cortex (PFC) and the ratings on the components of rapport modulated by prior synchronous/asynchronous physical movements.**

Three components of rapport: (a) mutual attentiveness, (b) positivity, and (c) coordination. Coloured points (•, ×) represent each dyad with the two conditions. Coloured lines represents the best linear fit for each dyad estimated by the repeated measures correlation, using the same slope and varying intercepts[1]

**Supplementary Text**

**Temporal independence of IBS from shallow signal synchronization**

To further confirm the neural origin of the observed IBS, we investigated whether the temporal changes in IBS were similar to the interpersonal synchronization of the shallow signals, which is expected to reflect synchronization in the physiological (e.g. arousal) and/or physical (head motion and postural) changes[2]. Temporal correlations between the interpersonal WTC time courses of brain signals obtained by using dual-SD regression[3] and the interpersonal WTC time courses of shallow signals from the 1-cm-SD pairs, at Fourier period 45 seconds, were calculated at each channel (left lateral and medial). The correlations were averaged over the two sessions, Fisher z-transformed, and subjected to a one sample t test. There were no significant correlations for the left lateral and medial channels (left lateral: mean $r = -0.05$, $t(31) = -0.62$, $p = 0.539$; medial: mean $r = 0.07$, $t(31) = 0.82$, p=0.419).

The same correlation analysis between the interpersonal WTC time courses of deep signal obtained from the 3-cm-SD pairs without using the dual-SD regression and the interpersonal WTC time courses of shallow signals showed significant correlations (left lateral: mean $r = 0.40$, $t(31) = 6.60$, $p = 2.2 \times 10^{-7}$; medial: mean $r = 0.43$, $t(31) = 5.38$, $p = 7.2 \times 10^{-6}$).

These results indicate that the IBS was independent from the artefactual interpersonal synchronization in the shallow tissues, while the IBS obtained without artefact removal using the dual SD regression was moderately contaminated by the artefactual synchronization in the shallow tissues.

### References


1 Bakdash, J. Z. & Marusich, L. R. Repeated measures correlation. *Front Psychol* **8**, 456 (2017).

2 Nozawa, T., Sasaki, Y., Sakaki, K., Yokoyama, R. & Kawashima, R. Interpersonal frontopolar neural synchronization in group communication: An exploration toward fNIRS hyperscanning of natural interactions. *Neuroimage* **133**, 484–497 (2016).

3 Saager, R. B., Telleri, N. L. & Berger, A. J. Two-detector corrected near infrared spectroscopy (C-NIRS) detects hemodynamic activation responses more robustly than single-detector NIRS. *Neuroimage* **55**, 1679–1685 (2011).
